# MicroRNA miR-219 is required for neural border and neural crest development in *Xenopus* neurulas

**DOI:** 10.64898/2026.06.09.730798

**Authors:** Alice M. Godden, Nicole Ward, Méghane Sittewelle, Rafeeq Mir, Aleksandr Kotov, Marco Antonaci, Anne H. Monsoro-Burq, Grant N. Wheeler

## Abstract

Neural crest (NC) multipotent stem cells give rise to many tissues including most of the peripheral nervous system, pigment cells and the craniofacial mesenchyme and skeleton. During gastrulation and early neurulation, cranial NC cells are specified in the ectoderm territory located between the anterior neural plate ectoderm and the future pre-placodal and lateral non-neural ectoderm. At the end of neurulation, NC cells undergo an epithelial-to-mesenchymal transition and migrate to various locations in the developing embryo where they differentiate. While the fine-tuning of NC specification is increasingly being elucidated, many questions remain, including how microRNAs may govern expression of gene programs during these processes. MicroRNAs are short non-coding 20-22 nucleotides-long RNAs which regulate gene expression through post-transcriptional repression. We have identified miR-219 as a candidate regulator of *Xenopus* NC development. Here, miR-219-dependent molecular pathways were investigated by morpholino knock-down and reveal NC phenotypes. The development of the NC and adjacent ectoderm was evaluated using whole mount *in situ* hybridization of key markers (*pax3*, *zic1, xhe2*, *sox10*, *snai2, sox2*), alcian blue cartilage staining, phenotype analysis, RNA sequencing of microdissected dorsal ectoderm and microRNA rescue experiments. While neural induction is mainly unaffected, miR-219 depletion alters gene expression programs associated with neural border development, resulting in loss of NC specification.

**Highlights:**

- miR-219 depletion expands the neural border territory and disrupts neural crest specification.
- miR-219 depletion phenotypes are rescued with miRNA mimics.
- miR-219 morphant neural border expansion is rescued by *pax3* depletion.
- RNA-seq reveals specific gene program modulation in miR-219 morphant neural crest.
- miR-219 is predicted to directly downregulate the neural gene *Hes5.3*.

## INTRODUCTION

The neural crest (NC) is a multipotent embryonic cell population that gives rise to a diverse array of cell types and tissues in the body (Alkobtawi and Monsoro-Burq, 2020). After induction in the dorsal ectoderm adjacent to the prospective neural plate, NC cells undergo an epithelial-to-mesenchymal transition and migrate away from the dorsal ectoderm, forming distinct migratory streams all along the body anterior-posterior axis. At the end of migration, NC cells differentiate into more than 30 different cell types, including components of the craniofacial skeleton, adrenal medulla, pigment cells, and various elements of the peripheral and autonomous nervous system (Aoto et al., 2015; Cheung and Briscoe, 2003). Specifically, cranial NC cells originate from the anterior part of the neural border (NB), a region of the dorsal ectoderm located on the side of the neural plate (NP), which also generates the cranial placodes. Cranial NC contribute to the formation of pigment, head mesenchyme, skull and facial skeleton, and cranial nerves and glial cells (Cordero et al., 2011; Gilmour et al., 2002; Simoes-Costa and Bronner, 2015). The broad developmental potential of neural crest cells also makes them particularly vulnerable to developmental dysregulation. Hence, neurocristopathies are a diverse group of congenital disorders resulting from defects in NC development or tumorigenesis, reflecting the wide range of cell types derived from the NC progenitors. These disorders include DiGeorge syndrome, Waardenburg syndrome, craniofrontonasal dysplasia (cleft palate), and cancers such as neuroblastoma and melanoma (Gouignard et al., 2016; Ward et al., 2018). Additionally, in amphibians, the dorsal ectoderm forms the hatching gland (HG) a tissue producing proteolytic enzymes allowing tadpoles to escape from their encapsulating vitelline membrane (Kurauchi et al., 2010). The HG also arises from the NB, along with the PPE (Hong and Saint-Jeannet, 2007).

MicroRNAs (miRNAs) are short, non-coding single-stranded RNAs, with a mature form of 20–22 nucleotides (Alberti and Cochella, 2017; Lee et al., 1993; Shah et al., 2017), modulating post-transcriptional gene expression. MiRNAs are located throughout the genome, and can be produced from intronic regions or exonic regions as independent genes from the genome (Olena and Patton, 2010). MiRNAs have been found across the animal kingdom, with many orthologues (Bartel, 2004; Godden and Immler, 2022; Godden et al., 2025a). In development, miRNAs have roles in C. *elegans, Drosophila* (Chandra et al., 2017), chick, mouse, frog, fish and human (Ahmed et al., 2015; Goljanek-Whysall et al., 2014; Mok et al., 2017; Ward et al., 2018). Recent studies have demonstrated that miRNAs also play direct roles in NC development (Gessert et al., 2010; Godden et al., 2021; Godden et al., 2025b; Ward et al., 2018). We previously found that miR-219 is expressed in NC and neural tissue in *Xenopus* embryos (Godden et al., 2021; Ward et al., 2018) and that CRISPR-mediated knockout of miR-219 results in craniofacial phenotypes, suggesting a role in cranial NC development (Godden et al., 2021). In this study, we aim to understand the requirement of miR-219 for NC development and to elucidate the mechanisms by which miR-219 influences the development of *Xenopus* NC.

## RESULTS AND DISCUSSION

To investigate the function of miR-219 in NC development, we employed several approaches including morpholino knockdown, whole-mount *in situ* hybridisation, and RNA-seq. Morpholinos (MOs) were designed to knock down (KD) the mature miRNA via base complementarity (Flynt et al., 2017). Injections of a non-specific MO mimic (control mimic) and of a mismatch MO (miR-219 MM) served as controls. The effects of miR-219 unilateral depletion on cranial NC development were assessed using Alcian blue staining to evaluate craniofacial cartilage formation. In stage 45 tadpoles, miR-219 depletion resulted in pronounced craniofacial phenotypes, including reduced size and disorganisation of the branchial arches, as well as shifts in Meckel’s cartilage and the palatoquadrate compared to the uninjected contralateral side of the embryo (Fig. 1). These results are in line with the previous results observed with CRISPR-Cas9 KO of miR-219 in *X. tropicalis*, where miR-219 crispant tadpoles feature distorted craniofacial phenotypes indicative of disrupted NC (Godden et al., 2021).

**Figure 1.**
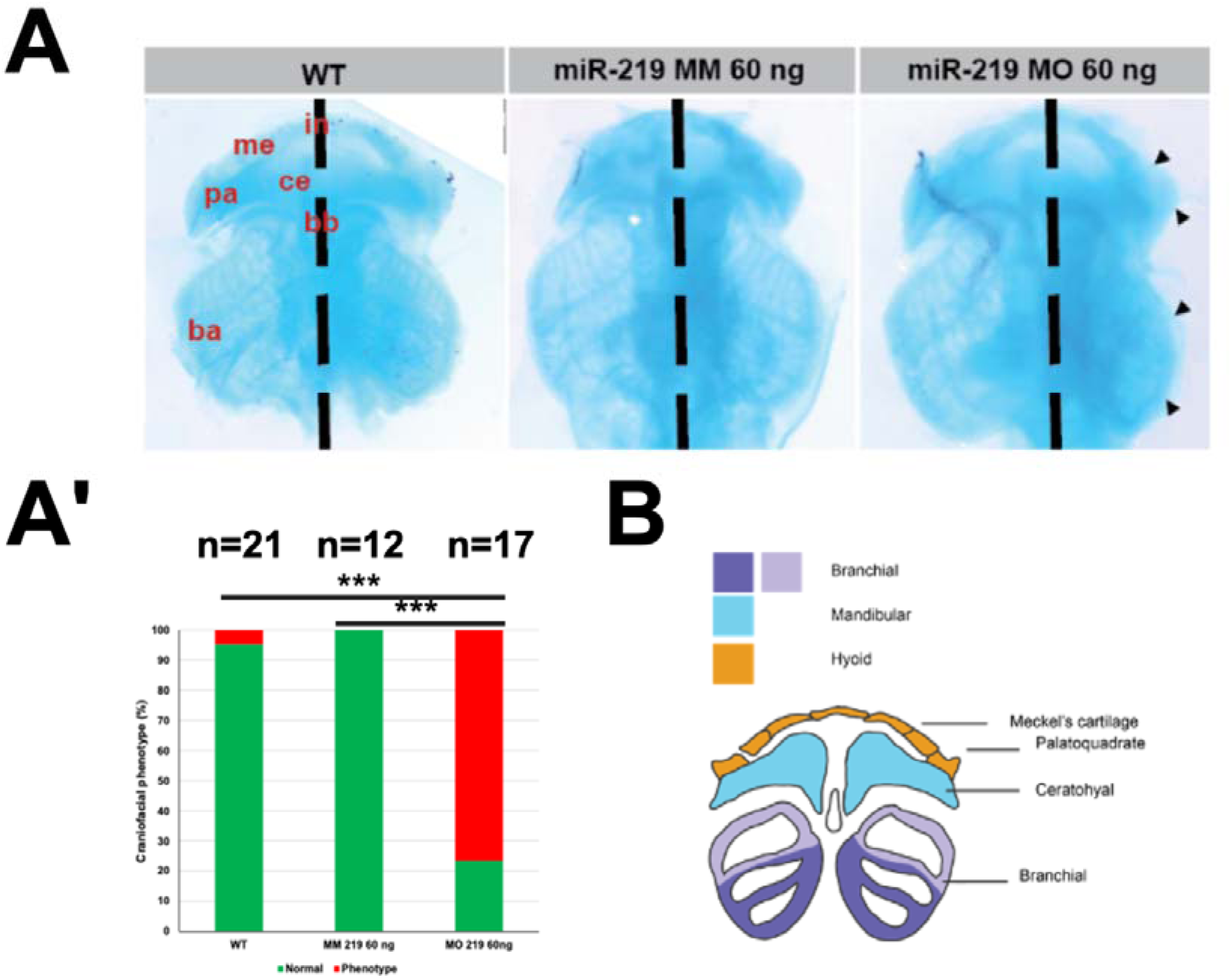
Assessment of craniofacial and craniofacial cartilage development after MO mediated miRNA KD. Embryos were injected into one dorsal blastomere at the 4-cell stage of development with a 10 nL calibrated needle and co-injected with 5 ng GFP cRNA. (A) Alcian blue cartilage preparations (st.45) show clear branchial arch and cartilage phenotypes following miRNA KD. (A’) Blind count phenotype data for alcian blue preparations. Test used for statistical analysis is Chi-squared on independent repeats. WT vs miR-219 MO p=<0.001, miR-219 MM vs MO p=<0.001. Key: ba= branchial arches, me= Meckel’s cartilage, pa= palatoquadrate, ce= ceratohyale, bb= basibranchiale, in= infrastrole. (B) Schematic of craniofacial cartilage in developing *Xenopus* tadpole at st. 45.

To further investigate the role of miR-219 during *Xenopus* dorsal ectoderm patterning, we first performed whole-mount *in situ* hybridisation on miR-219 morphant embryos after unilateral depletion (lineage-traced with betagalactosidase-derived red staining), using a well-characterized suite of genes marking each structure derived from the dorsal ectoderm (Hong and Saint-Jeannet 2007; Plouhinec et al, 2017): the neural plate (NP, *sox2*), the neural border (NB, *pax3* labels the NB including NC, placode and hatching gland (HG) progenitors, *zic1* also labels the NB and part of the lateral neural plate), the specified NC (*snai2* labels early specified NC in gastrula and neurula stage embryos*; sox10* labels late neurula-stage NC), and the specified HG (*xhe2*), (Fig. 2). MiR-219 depletion resulted in a marked reduction of specified NC and HG marker expression (Fig. 2Ad, k, w), while it expanded expression of the NB progenitor domain markers *pax3* and *zic1* (Fig. 2Aq; Fig. 3). Quantification of these results is shown in Figures 2B and 3B. In addition to affecting early NC specification, miR-219 MO knockdown also decreased expression of *twist1* in the migratory-stage cranial NC at tailbud stages. At that stage, the few NC cells that retained some *twist1* expression exhibited markedly decreased migration streams (SFig. 2). These observations indicate that NC early patterning is disrupted when miR-219 is depleted, and that this phenotype is not compensated during later stages of development. The severely reduced craniofacial NC streams accounts for the head cartilage defects seen at tadpole stage. Furthermore, given the role of miR-219 in regulating neural plate border-associated factors, we next examined HG development as an additional neural border–derived tissue and tested *Xhe2* expression. Following miR-219 KD a strong reduction in expression of *Xhe2* was observed (Fig.2 Aw) which was rescued by the miR-219 mimic (Fig. 2Ay). In conclusion, all NB-derived fates were altered upon depletion of miR-219.

**Figure 2.**
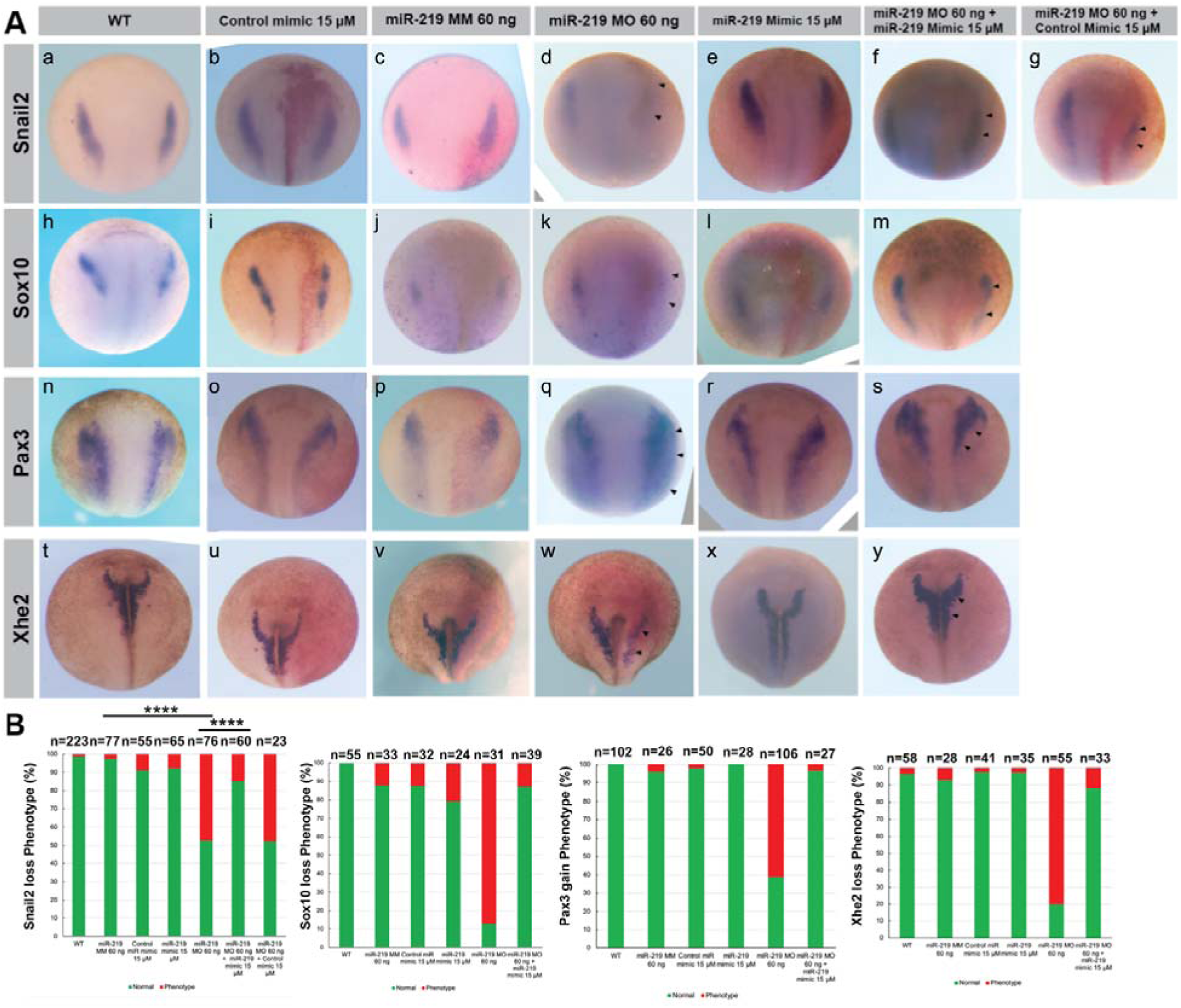
Functional characterisation of MO mediated miRNA KD of miR-219 with synthetic miRNA mimic rescues. Embryos were injected into one dorsal blastomere at the 4-cell stage of development with 300 pg of lacZ developed with Red-gal. (A) miR-219 MO KD and rescue for *Snail2*, (st.14), *Sox10*, (st. 14), *Pax3* (st. 14) and *Xhe2* (st.18) with control groups: Wild Type (WT), Control miR-mimic, miR-219 mismatch (MM), miR-219 MO, miR-219 mimic, miR-219 MO and miR-219 mimic and miR-219 MO and control miR-mimic. (B) Count data for phenotypes for each marker gene. *Snail2* was carried out with 3 individual biological experimental repeats, statistical analysis was performed using *Chi*-squared. 219 MM vs 219 MO *p* = 1.47 x 10^-10^, 219 MO vs Rescue *p* = 0.000054. Overall reduction in NC markers was seen along with deregulation of *Pax3* and *Xhe2* following miRNA KD. Relevant miRNA mimics successfully and specifically rescued phenotypes. MM= Mismatch control, MO= Morpholino.

**Figure 3.**
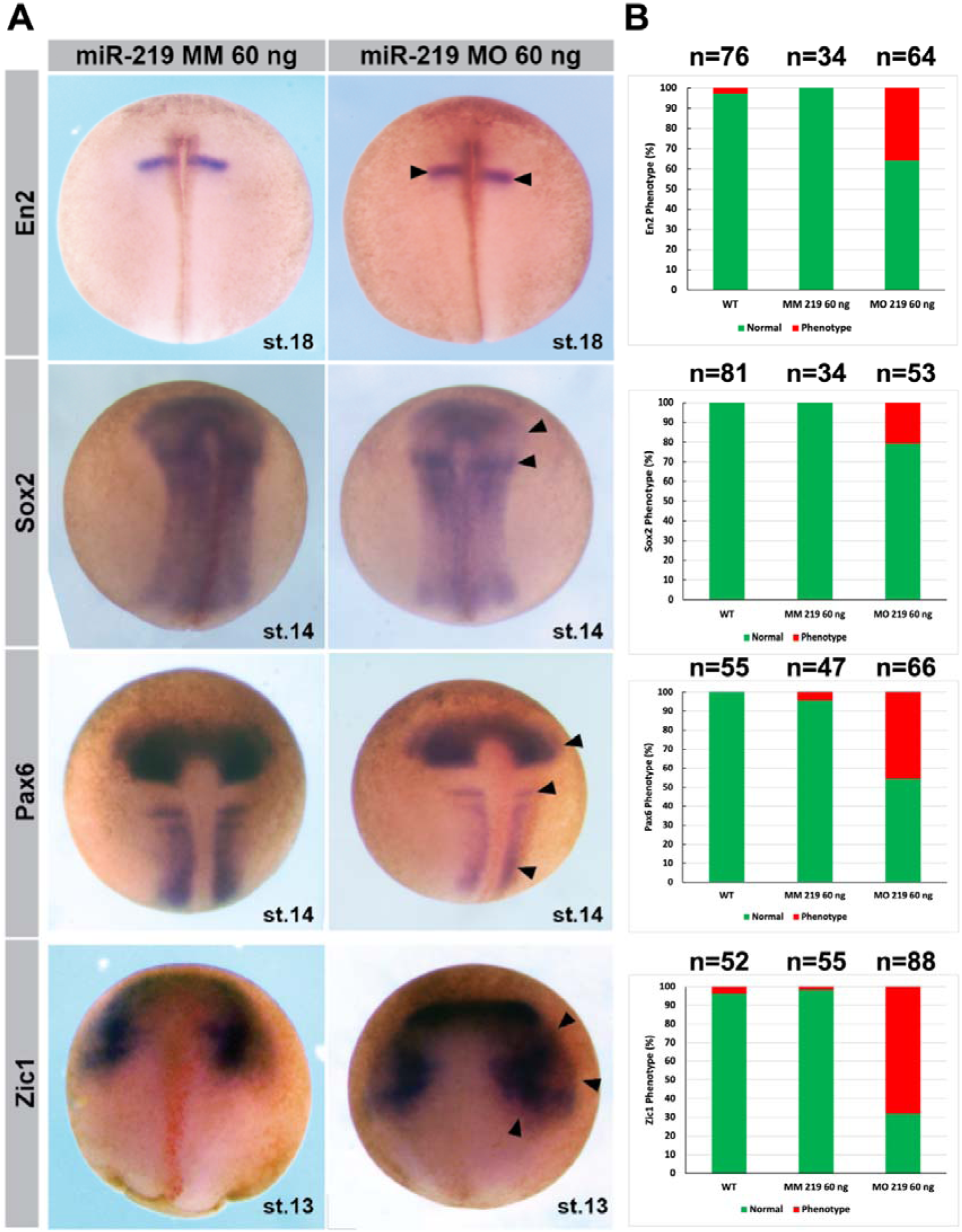
Assessment of neural, neural plate and placodal development following MO mediated miRNA-KD. Embryos were injected into one dorsal blastomere at the 4-cell stage of development with MO and 300 pg of lacZ developed with Red-gal. (A) Whole mount *in situ* hybridisation of *En2* (st. 18), *Sox2* (st. 14), *Pax6* (st. 14) and *Zic1* (st. 13) following MO mediated miRNA KD. (B) Blind score phenotype counts. MiR-219 MO KD exhibited a shift in anterior-posterior patterning with shifts in *En2*, *Sox2* and *Pax6* midbrain profiles and a subtle expansion of Zic1. MM= Mismatch control, MO= Morpholino.

To confirm the specificity of these phenotypes, we performed rescue experiments using synthetic miRNA mimics*. X. laevis* embryos were injected at the 4-cell stage into one dorsal blastomere with 300 pg *lacz* mRNA, MO, miRNA mimic (miR-219 mimic), or combinations thereof. No phenotypes were observed following injection with control miRNA mimic (cel-miR-39-3p), miR-219 mimic alone, or mismatch (MM) MO. Synthetic LNA miRNA mimics complementary to the mature miRNA sequence were used to rescue the phenotypes observed in whole-mount *in situ* hybridisation, as previously described for miR-196 KD (Godden et al., 2025b) SFig.1B). For all markers, the use of the relevant synthetic miRNA mimic successfully rescued the phenotypes generated by miRNA MO KD, while control mimics had no effect on embryo development. MiRNA expression levels were significantly increased by the mimic and decreased by the MO (SFig. 1); co-injection of mimic and MO restored expression levels of *snai2* (Fig. 2Af) and *sox10* (Fig. 2Am), while it reduced *pax3* expanded expression (Fig. 2As). These results demonstrate the specificity of the miR-219 KD effects (Fig. 2 and SFig. 1).

### Neural progenitor gene expression is also sensitive to miR-219 levels

Given that miR-219 is also expressed in the neuroectoderm (Godden et al., 2021) and that its knockdown affects neural plate border specification and Pax3-dependent patterning, we next examined whether miR-219 also influences broader neural and anterior–posterior axial patterning (Fig. 3). The engrailed gene family comprises highly conserved homeobox genes, with *En2* serving as a marker expressed at the midbrain-hindbrain boundary (Dur et al., 2020; Hemmati-Brivanlou et al., 1991). *En2* is a direct target of the Wnt signalling pathway and interacts with BMP antagonists and retinoic acid signaling during axial patterning of the *Xenopus* nervous system (Lou et al., 2006; McGrew et al., 1999)(Chen et al., 2001). MiR-219 KD resulted in a slight posterior shift of En2 expression without a clear change in expression intensity (Fig. 3). This indicates that the AP pattern is slightly modified, with a marginal anteriorisation of the neural plate.

*Sox2* marks the specified neural plate, while *zic1* labels the edges of the neural plate and the adjacent neural border territory (Fig. 3). After miR-219 depletion, *sox2* expression was mostly normal, with the slight AP shift observed with *en2* (Fig. 3).This suggested that the global neural induction occurred normally in morphant embryos. In contrast*, zic1* expression was expanded and upregulated around the neural plate following miR-219 depletion, as observed for *pax3* (Fig. 3). This finding confirms the unbalanced expression of critical genes specifiying the neural border territory (Hong and Saint-Jeannet, 2007), providing a potential mechanistic link between this altered patterning and NC phenotypes observed above.

### Micro-bulk RNAseq details miR-219-dependent programs in NB development

In order to identify miR-219-dependent gene programs during the early specification of NC cells, NB ectoderm was micro-dissected at stage 14 from embryos injected with miR-219 MO or control MM MO. The ectodermal layers were isolated, the underlying mesoderm removed, and the accuracy of the dissection was confirmed by probing donor embryos with *pax3* (for neural border) and *snai2* (for NC) as previously described (Godden et al., 2025b). Bulk RNA-sequencing was done on single explants dissected in triplicates (control MM) and quadruplicate (miR-219 MO). Principal component analysis (PCA) and correlation analysis showed that samples clustered according to MO treatment except for one outlier, a morphant sample removed from downstream analyses as per standard procedures (Fig. 4A, D).

**Figure 4.**
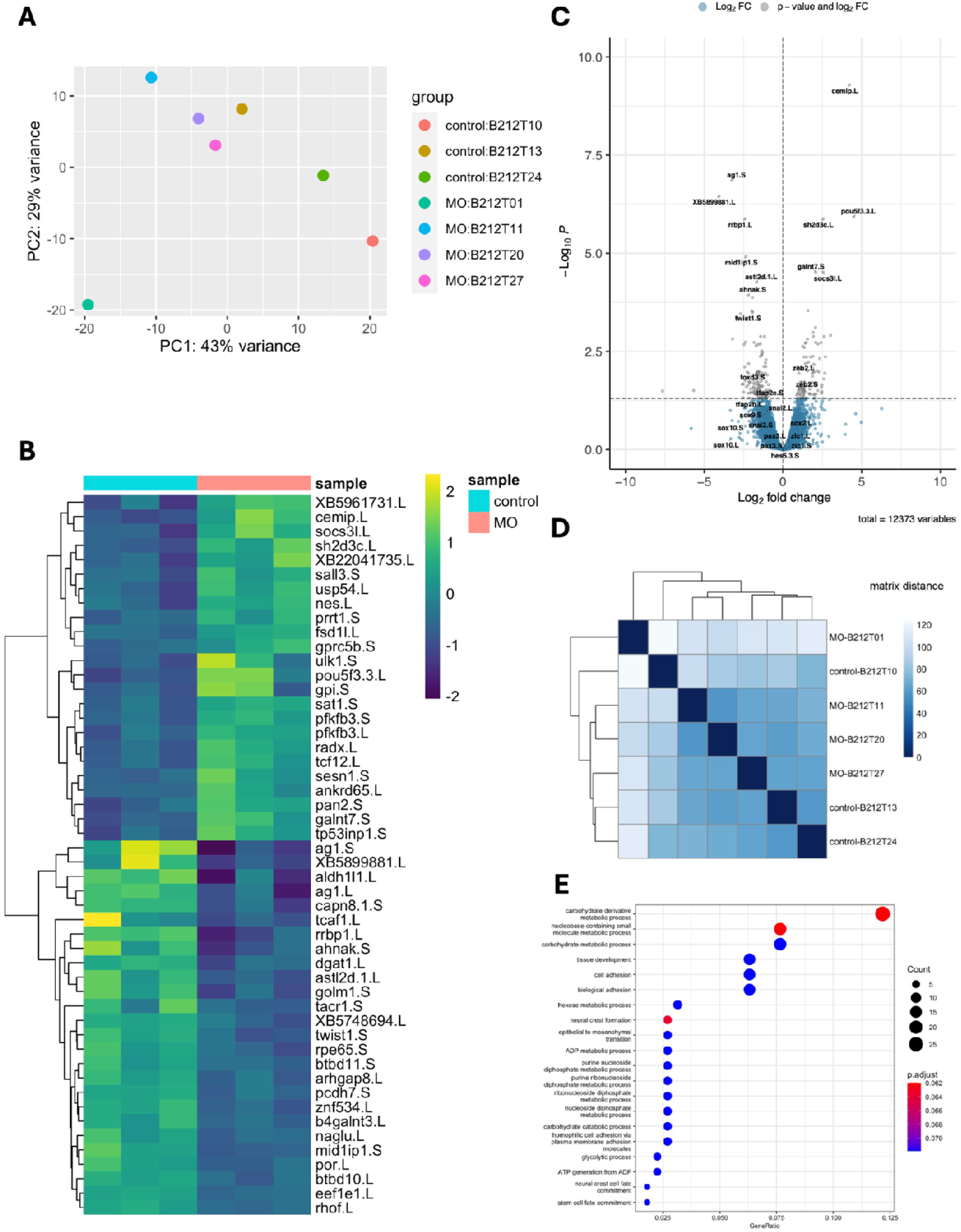
Bioinformatic analysis of gene expression following miR-219 KD. (A) Principal components analysis of sample clustering, with samples broadly clustering by treatment with miR-219 MO KD or miR-219 MM MO treatment at st.14. (B) Heatmap of significantly differentially expressed genes, clustered by treatment. (C) Volcano plot showing significantly differentially expressed genes with key NC, NPB, NP, HG and NNE, genes of interest labelled. (D) Clustering analysis of samples. (E) Top 20 GO terms analysis of significantly differentially expressed genes. The p-adjusted values for these terms are in colour and number of genes is noted by bubble size.

Heatmaps and volcano plots identified significantly differential gene expressions (Fig. 4B–C and SFig. 5), with strong reductions in NC markers such as *foxD3*, *twist1, pcdh8l,* and *pdgfra*. Other NC markers *(e.g. sox9*, *sox10* and *snai2)* were decreased below the logfold2 threshold following miR-219 KD. HG marker *xhe2* was also decreased. Genes that were upregulated are more likely to be direct miRNA targets, as miRNAs typically bind to complementary 3’ untranslated regions and post-transcriptionally repress gene expression (Rooda et al., 2020). The neural pluripotent progenitor markers *pou5f3.3* (also called *pou60)* (Young et al., 2014) and *sall3* (Deimling and Drysdale, 2009) were significantly upregulated, neural progenitor genes *sh2d3c* and *pfkfb3* were also more expressed (Fig. 4B-C)(Giudetti et al., 2014; Pegoraro et al., 2013). This suggested that the dissected tissue, located at the position of the NB, exhibited increased expression of immature neural progenitors. Neural and NB progenitor marker *zeb2* (van Grunsven et al., 2006), was also increased, as were NB markers *pax3* and *zic1*, albeit below the logfold2 threshold. Some markers shared between neural ectoderm and placode (*pax6, prdm12)* were also enriched (SFig. 5). Gene ontology (GO) analysis of significantly differentially expressed genes showed enrichment for terms related to NC formation, cell fate commitment, metabolic processes, and stem cell fate commitment (Fig. 3E). Last, the enrichment of GO terms associated with cell fate commitment, metabolic processes and NC formation, supported by increased expression of *pfkfb3 .L* and *.s* homeologs, was consistent with previous studies showing roles for some glycolysis-related metabolism regulators upstream of the fate decisions patterning the dorsal ectoderm (Pegoraro et al., 2015). In sum, the RNA-seq analysis of the cells formed in the morphant NB domain revealed a severe mispatterning of the NC program (*foxD3*, *twist1, pcdh8l, pdgfra)* with lower differential expression of some genes which were confirmed by WISH (*sox10, snai2)*, in parallel with the upregulation of genes associated with immature neural plate fate (*pou5f3.3, sall3)*, immature NB fate (*pax3, zic1, zeb2)* and some placode markers.

**Figure 5.**
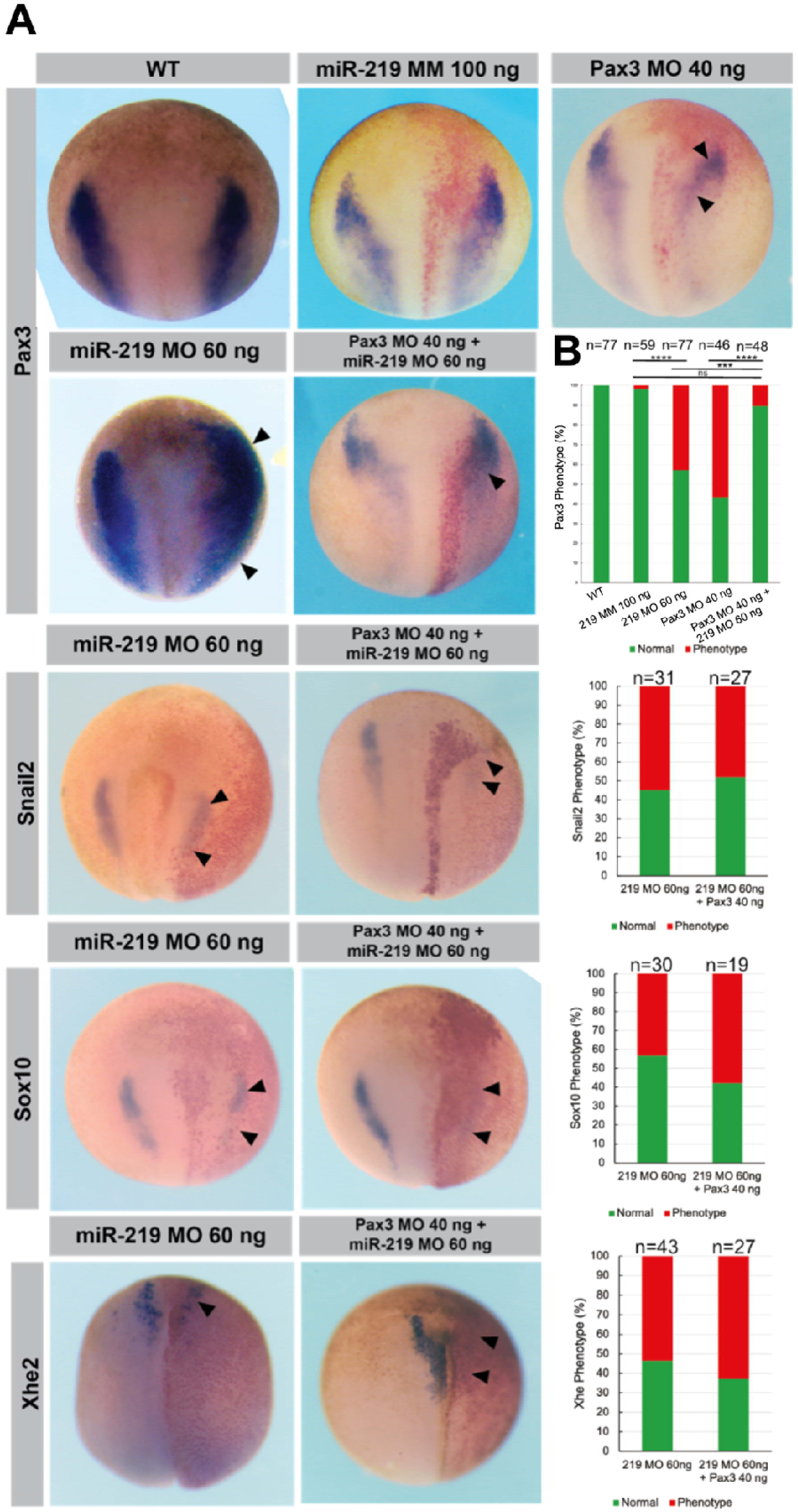
Rescuing miR-219 phenotypes with Pax3 MO. Embryos were double injected, the total dose of MO is indicated. Embryos were injected at the 2-cell stage of development into one blastomere, twice. 300 pg of lacZ was used as a tracer and developed with Red-gal. (A) *Pax3* (st. 14), *Xhe2* (st. 18), *Sox10* (st. 14) and *Snail2* (st. 14) whole-mount *in situ* hybridisation following MO KD with miR-219 MM MO, miR-219 MO, Pax3 MO, miR-219 MO with Pax3 MO or WT control. (B) Blind score phenotype counts. For Pax3 group three individual biological experiments were carried out, Chi-squared statistical testing was performed. 219 MM vs 219 MO p= 3.9 x 10^-8^, 219 MM vs 219 MO + Pax3 MO p=0.051, 219 MO vs 219 MO + Pax3 MO p=0.00013, Pax3 MO vs 219 MO + Pax3 MO p= 2 x 10^-6^. miR-219 MO phenotype was only rescued with Pax3 MO for Pax3 group. MM= Mismatch control, MO= Morpholino.

*In vivo*, different possible cell fates are specified from the neural border progenitors by a delicate balance of regulator gene expressions. For instance, while moderate levels of *pax3* and *zic1* cooperate to induce the NC fate by directly activating *snai2* (Kotov et al., 2024; Monsoro-Burq et al., 2005), elevated *pax3* levels repress *snai2* expression and favor hatching gland fates (*xhe2)* while elevated *zic1* levels favors placode specification (Hong and Saint-Jeannet, 2007). Here, miR-219 depletion caused a marked expansion of the *pax3*-positive territory in the superficial ectoderm layer (Fig. 2Aq, SFig. 4B), which may contribute to the observed reduction in NC markers. Similarly, *zic1* expression is enhanced in the lateral NB ectoderm (Fig. 3). In addition, in the micro-dissected tissue located at the level of the wild-type NB, further ectoderm specification into NC seems blocked while progenitor gene expressions are sustained. Together, these findings suggest that miR-219 regulates an upstream program involved in the broader patterning of the dorsal ectoderm, limiting the preplacodal ectoderm (PPE) fates, and favoring the specification of a subset of neural border derivatives including NC and hatching gland progenitors. Such early program may act upstream of Wnt, FGF, or BMP signalling pathways to regulate *pax3* and *zic1* domain of expression (Alkobtawi et al., 2021; Pla and Monsoro-Burq, 2018).

### Validation of genes modulated by miR-219 depletion identified by RNA-seq

In support to this analysis, we conducted an *in silico* miRanda analysis (https://github.com/hacktrackgnulinux/miranda) which predicted that *snai2* (NC)*, eya1, six3, six4* (placode progenitors), and *hes5.3* and *otx2* (neural plate progenitors) were directly targeted by miR-219 (Table 3). First, we examined the early PPE marker *Eya1* using a Luciferase reporter assay on chicken fibroblasts which validated that *eya1* is a direct target of miR-219 (SFig. 6). Next, we probed for *hes5.3* in whole embryos and find that its expression is strongly enhanced upon miR-219 depletion (SFig. 7). Thus, we find that miRanda predictions are validated either in a cellular model or in vivo, therefore supporting our hypothesis that miR-219 plays a role in general dorsal ectoderm patterning upstream of NC - PPE - NP fate choices.

**Figure 6.**
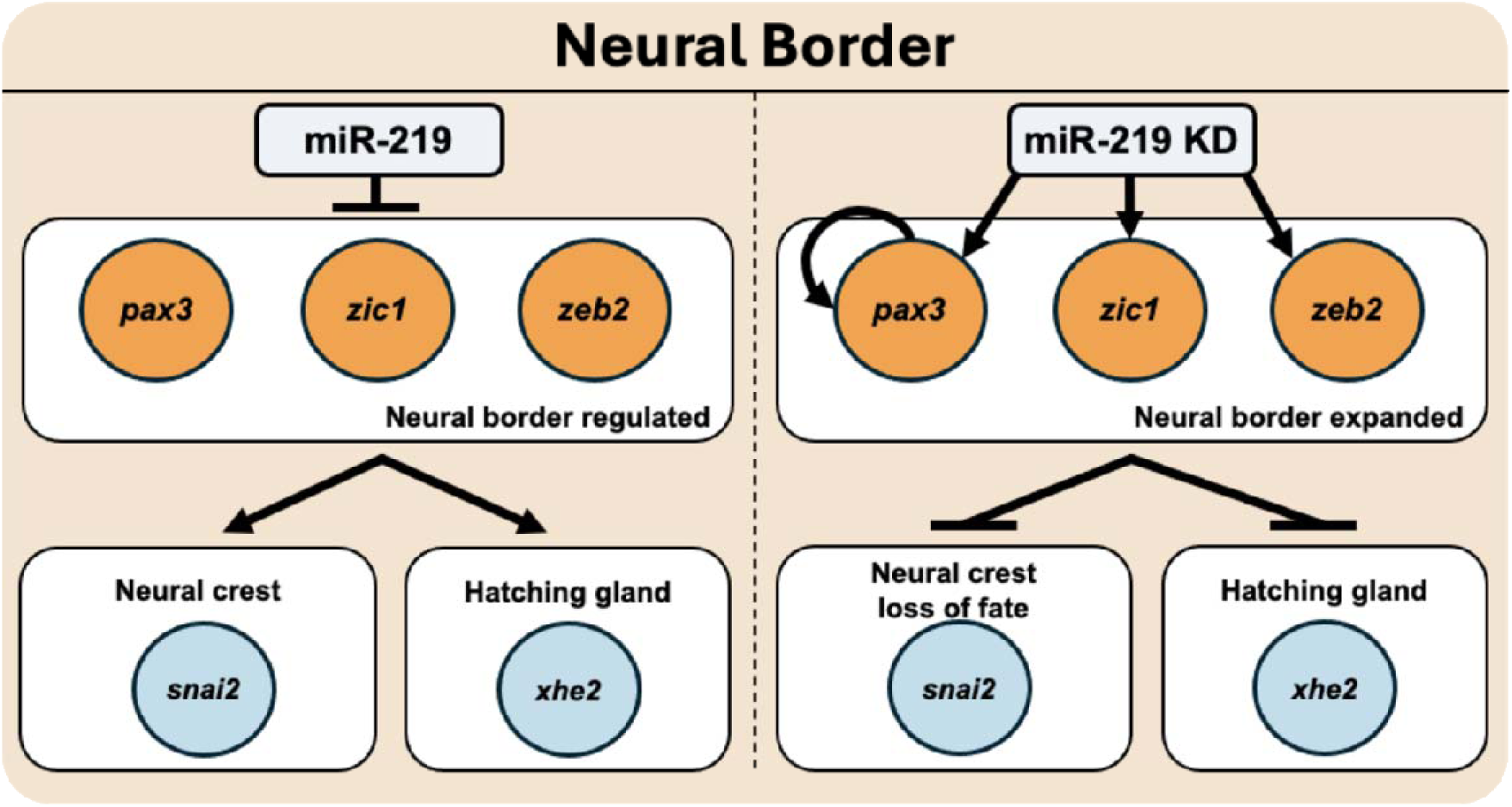
Model of the impact of miR-219 KD on the neural border. Blue circles indicate experimental data to show depletion of that gene, and orange colour circles represent enrichment of that gene. Mir-219 helps define the neural border leading to specification of the neural crest and hatching gland .

Taken together, the broad patterning defects observed following miR-219 knockdown suggest that miR-219 exerts its effects through regulation of multiple downstream targets. Consistent with this, miR-219 is predicted to target several genes relevant for dorsal ectoderm early patterning, including *snai2* and *hes5.3* (see Table 3 for miRanda analysis of miRNA target genes). Both genes were differentially expressed in RNA-seq data following morpholino-mediated knockdown of miR-219 (Fig. 4) and validated *in vivo* (Fig. 2 for *snai2)*: following miR-219 KD, *hes5.3* expression on the injected side of the embryo appeared increased expression in neural regions (Fig. S7), further supporting a role for miR-219 in regulating neural patterning and its downstream specification. These roles in key neural and NC specifiers suggest that miR-219 plays multiple roles during the initial step of each tissue induction and specification.

### Restoring lower pax3 levels is not sufficient to revert the miR-219 morphant phenotype

Given the striking expansion of *pax3* expression and the neural border defects observed on multiple other NB genes (*zic1, zeb2)* following miR-219 knockdown, we next examined whether *pax3* contributes functionally to these phenotypes. We used a validated pax3 MO (Monsoro-Burq et al., 2005) to attempt rescue of the expanded *pax3* expression domain in miR-219 morphants. *pax3* MO dose-response effects on *pax3* and *snai2* expression and cryosectioning of embryos from these experiments are shown in SFigs. 3-4. While miR-219 MO injections increased *pax3* domain of expression laterally, towards the PPE and non-neural ectoderm areas, *pax3* MO co-injections with miR-219 MO reduced *pax3* expression back to its normal spatial extent (Fig. 2Aq; Fig. 5; Fig. S4). This indicated that if the PAX3 protein cannot be synthesized (upon *pax3* MO blockade), *pax3* gene expansion does not occur even when miR-219 is depleted. PAX3 has been shown to autoregulate its own expression during NC development (Plouhinec et al., 2014). Moreover, neural border patterning involves a positive feedback loop involving PAX3, ZIC1, HES4 and TFAP2A (de Croze et al., 2011). Theoretically, such autoregulation may create an unlimited expansion of *pax3* expression and NB territory, unless a regulatory mechanism counteracts this activity. Our result suggests that one of miR-219 roles in NB patterning is to limit the *pax3/zic1-*positive domain and prevent its ectopic expression towards the non-neural ectoderm area.

We next evaluated if the interruption of *pax3* expansion would be sufficient to also restore the morphant phenotype of the other markers, and therefore revert the mispatterning phenotype. We examined the expression of *xhe2*, *sox10*, and *snai2* (Fig. 5). As before, the three markers were reduced following miR-219 MO knockdown (Fig. 2A,B; Fig. 5). However, co-injection of Pax3 MO led to a more frequent reduction in *sox10* and *xhe2* expression, while *snai2* levels remained similarly reduced (Fig. 5). This result indicates that restoring the extent of *pax3* domain of expression is not sufficient to restore the wildtype phenotype for the other NB markers. Rather NB patterning remains very disrupted. This suggests several mechanistic possibilities. The first hypothesis is that although the domain of expression of *pax3* seems rescued, the use of *pax3* MO prevents translation of the PAX3 protein. As PAX3 protein is essential within the NB, for further patterning of the different fates, NB formation would remain blocked. Additionally, the second hypothesis is that *pax3* alone is not sufficient to compensate the effects of miR-219 depletion, and that additional players would need to be re-adjusted to rescue the morphant phenotype. Such players may be, for example, the gene *zic1*, another essential NB specifier. These results suggest that miR-219 acts on multiple early steps of neural border specification.

In conclusion, miR-219 depletion and rescue experiments show that miR-219 is required for neural plate border patterning and neural crest development in Xenopus. Its loss does not affect the extent of the neural plate as shown by sox2 expression but expands pax3 and zic1 expression at the neural border, reducing neural crest specification markers, and leading to craniofacial cartilage defects. We propose that miR-219 acts as a post-transcriptional modulator that constrains neural border output and enables successful neural crest specification.

## MATERIALS AND METHODS

### Ethics

All experiments were carried out in accordance with relevant laws and institutional guidelines at the University of East Anglia, with full ethical review and approval, compliant to UK Home Office regulations.

### Embryo generation and microinjection

To obtain *X. laevis* embryos, females were primed with 100 units of PMSG and induced with 500 units of human chorionic gonadotrophin. Eggs were collected manually and fertilised *in vitro*. Embryos were de-jellied in 2% L-cysteine, incubated at 18°C and microinjected in 3% Ficoll into 1 cell at the 2-cell stage in the animal pole with 5 nL of enhancer reporter plasmid at 400 ng/μL or GFP capped RNA as control. Embryos were left to develop at 23°C. Embryo staging is according to Nieuwkoop and Faber normal table of *Xenopus* development (Gordon et al., 1994). GFP/LacZ capped RNA for injections was prepared using the SP6 mMESSAGE mMACHINE kit, 50 pg was injected per embryo.

Embryos were injected using a 10 nL calibrated needle. MO dose was optimized to 60 ng for miRNAs; MO and lacZ were injected at 4-cell stage of embryo development into the right dorsal blastomere. MiRCURY LNA miRNA mimics were used to replace miRNA in MO miRNA KD rescue. miR-219 mimic was used from (Qiagen, 339173 YM0047076-ADA, MIMAT0000276); hsa-miR-219a-5p miRCURY LNA miRNA Mimic, compatible with xtr-miR-219 sequence: 5’UGAUUGUCCAAACGCAAUUCU. A negative control miRNA mimic recommended by Qiagen was used (Qiagen, 331973 YM00479902-ADA); Negative control (cel-miR-39-3p), sequence 5’UCACCGGGUGUAAAUCAGCUUG.

To rescue expanded Pax3 phenotypes Pax3 MO (Table 1) was optimized at 40 ng to provide a reduction in Pax3 expression. This was co-injected with miR-219 MO 60 ng. As this isn’t possible in one injection MOs were made up so Pax3 MO final concentration was 20 ng and miR-219 MO was 30 ng, two injections into the embryo at 4 cell stage into 2 blastomeres then gave a concentration of Pax3 MO 40 ng and miR-219 MO 60 ng. For miR-219 MO MM two injections of 50 ng gave a final dose of 100 ng MO.

**Table 1.**
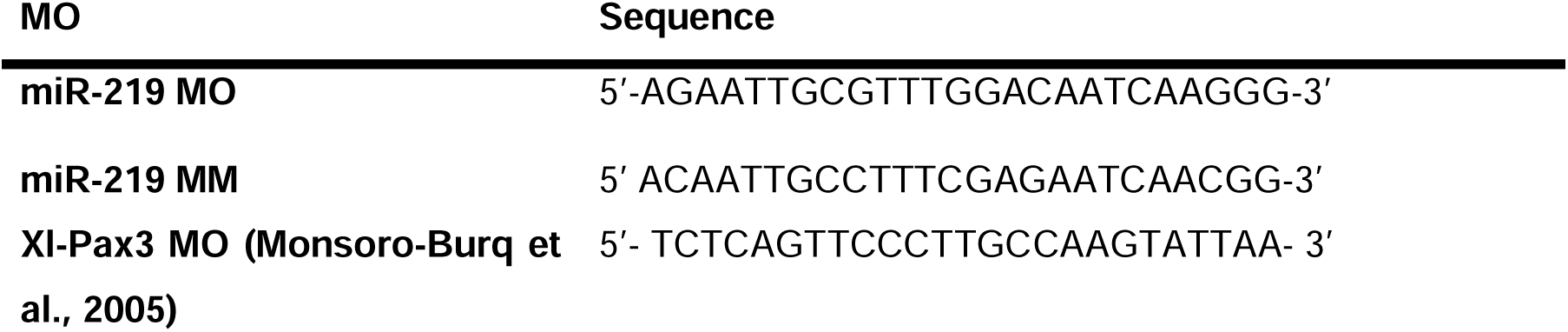
Injected MO sequence data.

*Chi*-squared test for association was used to test phenotype yes or no categories for MO injected embryos to see if there was a relationship between two categorical values. Excel was used to collate and tabulate data. IBM SPSS v25 to carry out Chi-squared test. When describing statistical significance; *p* <0.05 = *, *p* <0.01 = **, *p* <0.001 = ***, *p* =<0.0001= ****. Embryos were frozen on dry ice before RNA extraction. Total RNA was extracted from five st.14 *X. tropicalis* embryos, embryos were homogenised with a micro pestle, and RNA was extracted according to manufacturer’s guidance, Quick-RNA Mini prep plus kit (Zymo, Cat no. R1058). Samples were eluted in 25 µL of nuclease free water; RNA concentration and purity quantified on a Nanodrop 1000 and 1 µL was checked on a 2% agarose gel.

To produce cDNA for q-RT-PCR, miRCURY LNA RT kit (Qiagen, Cat No./ID: 339340). 50 ng of RNA was used and kit according to manufacturer’s instructions. cDNA was produced on a thermocycler with the following programme: 42°C for 60 min and 95°C for 5 min. cDNA was diluted 1:40 for q-RT-PCR. cDNA can be stored at -20°C. qRT-PCR reactions were set up in 10 µL volume containing 4 µL cDNA, 1 µL primer (10 µM for standard oligo primers, in accordance with manufacturer dilute for Qiagen LNA miRNA primer), and 5 µL SybrGreen (Applied Biosystems 4309155). See Table 2 for primer sequences and details.

**Table 2.**
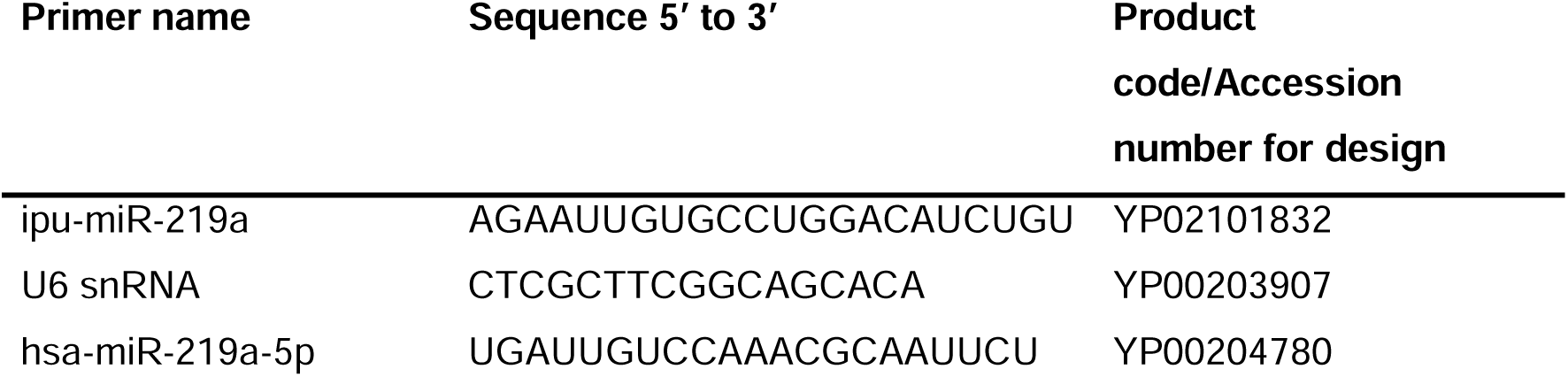
q-RT PCR Primers used for *X. tropicalis* embryos. miRCURY LNA miRNA PCR primers, Qiagen. mRNA primers were ordered as standard oligos.

**Table 3.**
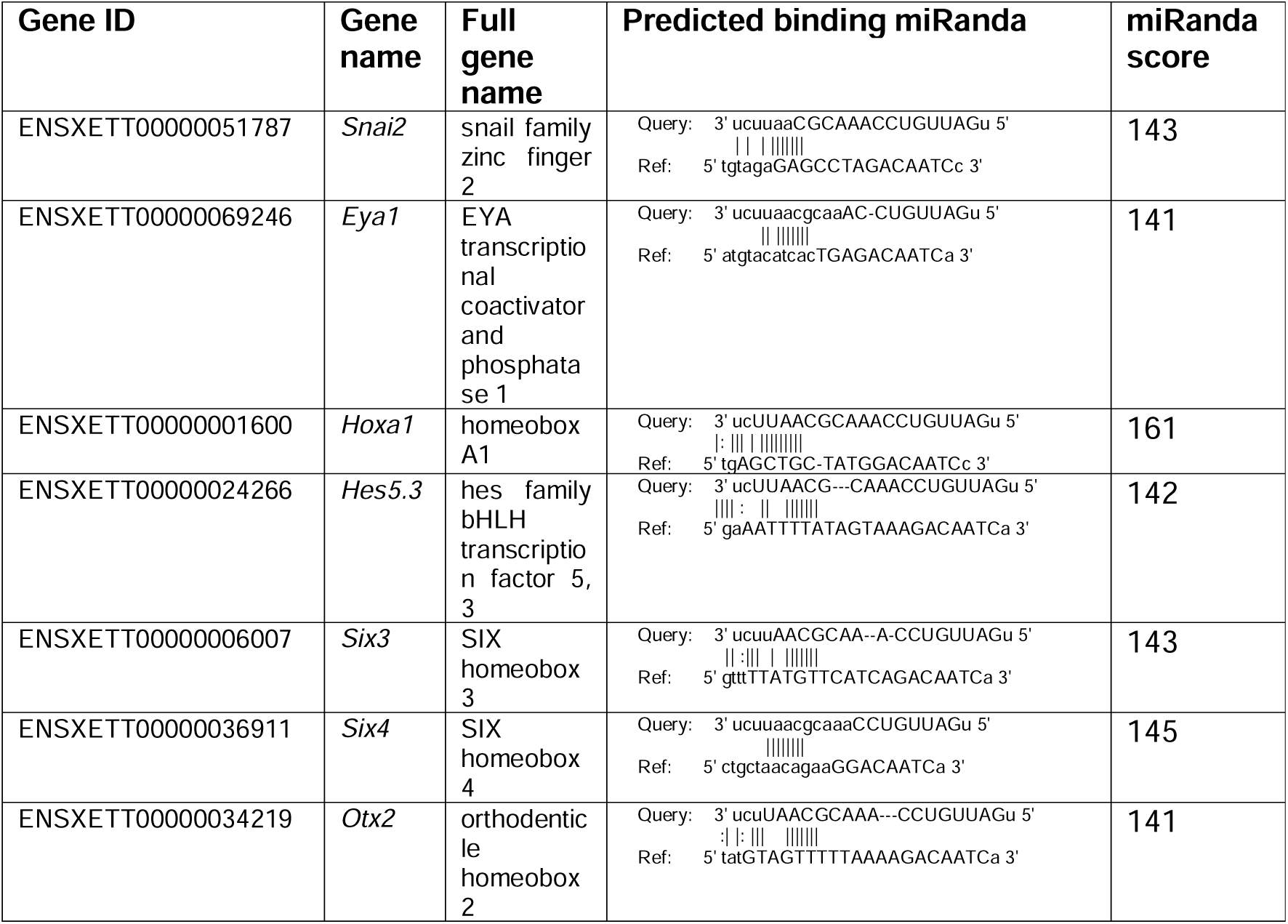
Results of miRanda analysis for miR-219. Contents cover Gene ID, name and predicted binding of miR-219 (query) and 3’ UTR of gene (Ref), with miRanda score in last column, higher score indicates stronger match.

WISH with LNA probes was carried out according to (Sweetman et al., 2006). Other in situs and probe synthesis were carried out according to (Monsoro-Burq, 2007; Sive et al., 2007). Plasmid information for generation of riboprobes can be found in Supplementary Table 1.

For RNA-sequencing embryos were injected into one blastomere at 4 cell stage with one of four MO’s (miR-219, miR-219MM) into one dorsal blastomere to target neural and NC tissue in one side of the embryo only. Embryos were left to develop until stage 14. One group underwent WISH to check NC genes were knocked down (data not shown) and the other group underwent NC dissections and RNA was extracted. Three replicates were collected for each condition (MO, MM and non-injected control). RNA samples underwent quality control using a Bioanalyser and q-RT-PCR was used to further validate the KD of NC specific genes in MO injected samples. Samples were then processed to Illumina sequencing.

### Luciferase assay

A Luciferase sensor construct for the EYA1-3’UTR was generated using sequences originating from chicken tissues, however EYA1 is homologues to the *Xenopus tropicalis* variant (SFig. 6). Mutant pGL3-EYA1 construct pGL3 mutant constructs were generated using the FastCloning strategy developed by (Li et al., 2011). A pair of primers placed in the ampicillin resistance gene were used as universal primers in the mutagenesis cloning. Primers were designed over the miRNA target site which contained mismatched nucleotides chosen to create an enzyme restriction site out of the miRNA target site (SFig. 6A-B). The two halves of each pGL3 construct were amplified using Phusion High Fidelity DNA polymerase (Finnzymes, NEB). Phusion PCR: 2.5 µL Forward primer (10 µM), 2.5 µL Reverse primer (10um) and 20ng pDNA were combined in a PCR tube with the mastermix. Once finished, 5 U Dpn1 restriction enzyme (Promega) was added to the PCR mixture and incubated on a thermocycler at 37°C for 2 h to digest any remaining pDNA. PCR products were verified on an agarose gel. 1 µL of each the products were transformed into 200 µL of DH5α competent cells for recombination.

Chicken DF1 fibroblast cells were cultured in Dulbecco’s modified Eagle medium (DMEM) containing GlutaMAX, 1g/L D-glucose and pyruvate (Life Technologies), with 10% fetal bovine serum (FBS) (Gibco, Life Technologies) and 1% penicillin-streptomycin (Gibco, Life Technologies) added on first use. Cells were passaged every other day by treatment with 1ml 0.25% Trypsin-EDTA (Gibco, Life Technologies) for 30 secs at RT followed by removal of trypsin and further incubation at 37°C for approximately 2 mins. Cells were then transferred to a new flask containing media at a 1:4 dilution. Cells were maintained in a humidified cell culture incubator at 37°C with 5% CO2. 12.3.2. Cell transfection Cells were detached from the flask surfaces as described in 9.3.1 and counted using a haemocytometer. Approximately 7,000 cells were added to each well of a 96-well plate and media added to make a volume of 100 µL. Cells were incubated for 24 h to allow re-attachment to the plate surface. The medium was removed and replaced with 50 µL serum-free medium. A transfection mix was prepared as described below. Transfection mix 1: 100 ng pGL3 pDNA 50 nM miRNA mimic 25 ng Renilla pDNA x µL Serum free media to a total volume of 25 µL Transfection mix 2: 25 µL Serum-free media 0.2 µL Lipofectamine 2000 Transfection reagent (Life Technologies) 48 Tubes were left at RT for 5 mins. Transfection mixes 1 and 2 were combined and left at RT for 20 mins. 50 µL of the combined transfection mix was added to the wells followed by 50µL of serum-free media and cells were left to incubate at 37C for 24 h in a humidified incubator. Custom siRNA oligos (Sigma) were used as miRNA mimics. Each construct was evaluated in the presence of the putative target miRNA mimic, a control miRNA (MISSION siRNA Universal negative control (#1) from Sigma) and without a miRNA mimic. 12.3.3. Luciferase assay A Dual-Luciferase Reporter Assay System kit was used to assess luciferase activity (Promega). This kit allows Renilla Luciferase to serve as an internal control of transfection efficiency. Cells were washed twice with cold PBS and 60 µL 1x Passive Lysis buffer was added to each well and the plate was rocked gently for at least 15 mins. 10 µL lysis solution and 50 µL luciferase assay reagent II was combined, and photo emission was measured at 562 nm using a multi-label counter (Perkin Elmer). 50 µL Stop and Glo reagent was added and the Renilla Luciferase photo emission measured at 562nm. 12.3.4. Normalisation of luciferase assay data Three biological replicates, each with four technical replicates, were performed. Each firefly luciferase assay reading was normalized to its Renilla luciferase reading. The average activity of the plasmid transfected with control mimic was set to 100%. The activity of the constructs transfected with the putative target miRNA mimic was normalized to the average activity of the plasmid transfected with the control mimic. A Mann-Whitney test was performed using GraphPad Prism version 6 for Mac (GraphPad Software, San Diego California, USA, www.graphpad.com). Validation of Eya1 as a direct target was done by luciferase assays of modified pGL3 reporter constructs containing the 3’UTR of Eya1. This bioluminescence assay is a quantitative method based on sequential measurements of Firefly and Renilla luciferase activities in a single sample. The 3’UTR of Eya1 contains one conserved predicted miR-219 binding site which was mutated to show that repression is mediated specifically through the predicted target site. In summary, WT or mutant constructs (100ng) were co-transfected, into chicken DF1 fibroblast cells, with Renilla vector (25ng) and either with or without the miR-219 siRNA (si-219). This siRNA is used to mimic the action of miR-219. A universal negative control siRNA (siC; Sigma) was used as a negative control.

### Data analysis

RNA-sequencing reads were mapped to the *X. laevis* v10.1 genome assembly using STAR (v.2.7.3a), (Dobin et al., 2013). Differential expression analysis was carried out using DESeq2 (v.1.32.0), (Love et al., 2014) in R (v.4.1.1). Genes with an adjusted *p*-value below 0.05 and had a log2fold-change greater than 1 were considered significant and were reported by the workflow. The gene model used in the DE bioinformatic analysis was *X. laevis* (NCBI v10.1). For GO enrichment analysis of a DE genes we used ClusterProfiler (v.4.0.5). MiRanda was used to analyse miRNA-219 3’ UTR gene targets with default settings (John et al., 2004). Mature miRNA sequences were accessed from miRbase (Griffiths-Jones et al., 2006), and annotation and reference fasta file accessed from Ensembl v106 *Xenopus tropicalis* v9.1 genome miR-219 was not annotated in the *X. laevis* genome, but is highly conserved. RNA-seq data are available at the GEO database under accession: GSE289706.

For Alcian blue craniofacial cartilage staining embryos were dehydrated in ethanol and were then left in Alcian blue (20 mg) for 3 nights. After this they were washed 3 times for 15 minutes in 95% ethanol (Sigma, UK) then rehydrated in 2% KOH using 10 minute washes of 75% ethanol in 2% KOH, 50% ethanol in 2% KOH, 25% ethanol in 2% KOH then 3 x 2% KOH washes. Embryos were then stored in glycerol with 1 hour washes of 20% glycerol in 2%, 40% glycerol in 2% KOH, 60% glycerol in 2% KOH and finally stored in 80% glycerol in 2% KOH. Embryos were washed 3 x 5 mins in PBS before 20 min incubation in 10 mL pre-incubation solution (0.5 X SSC (150 mM NaCl, 15 mM sodium citrate, pH 7.2), 0.1% Tween. 20), and then embryos were incubated in 10 mL of depigmentation solution (5% formamide, 0.5 X SSC, 3% H2O2). Embryos were then carefully dissected under a microscope using fine forceps to remove outermost membranes surrounding craniofacial cartilage. Method for this clearing was based on (Affaticati et al., 2018).

Embryos were imaged on agarose dishes with a Zeiss Axiovert Stemi SV 11, Jenoptik ProgRes C5 camera (Germany), ProgRes software version 2.7.6. Fluorescent images were captured using Leica MZ 16 F microscope, Leica DFC300 FX camera, Leica Kubler codix light source, Leica FireCam software version 3.4.1.

## ACKNOWLEDGEMENTS

We would like to thank Prof. Jean-Pierre Saint-Jeannet and Dr. Maggie Walmsley for plasmids, Prof. Andrea Münsterberg and Prof. Simon Moxon for helpful discussions and the Wheeler, Münsterberg, and Monsoro-Burq labs for support. A Luciferase sensor construct for the EYA1-3’UTR was generated by Dr. Camille Viaut from the MuLJnsterberg lab and generously provided for this experiment.

This work was supported by the UKRI Biotechnology and Biological Sciences Research Council Norwich Research Park Biosciences Doctoral Training Partnership (Grant numbers BB/M011216/1 to GW/AG and BB/J014524/1 to GW/NW). This project also received funding from European Union Horizon 2020 Marie Skłodowska-Curie grant no. 860635, NEUcrest ITN (AK, MA, AHMB, GW); Agence Nationale pour la Recherche ANR-15-CE13-0012-01, ANR- 21-CE13-0028; Institut Universitaire de France (AHMB); Fondation pour la Recherche Médicale (AHMB). RNA sequencing used the ICGex NGS platform of Institut Curie (ANR-10-EQPX-03; ANR-10-INBS-09-08; Canceropole Ile-de-France; SiRIC-Curie INCa-DGOS-4654)

## AUTHORS CONTRIBUTIONS

GW and AMB designed the study and directed the project. AG, NW, MS, MA and AMB performed the experiments and acquired the data. AG and AK analysed the bioinformatics data. GW, AMB, NW and AG analysed the data. AG, GW and AMB and wrote the manuscript.

## Abbreviations

KD: Knockdown
MM: Mismatch Morpholino
MO: Morpholino
NC: Neural crest
NCP: Neurocristopathy
NNE: Non-neural ectoderm
HG: hatching gland
PPE: pre-placodal ectoderm

