## Supplemental Data and Figures for "MicroRNA miR-219 is required for neural border and neural crest development in *Xenopus* neurulas"

### Supplementary Data

Supplementary Table 1- Riboprobe synthesis plasmids information

| Clone name | Antibiotic resistance | Backbone | Sense RE | Sense polymerase | Antisense RE | Antisense Polymerase | Insert Size | Source |
|---|---|---|---|---|---|---|---|---|
| Pax3 | Ampicillin | pBSK | | | BglII | SP6 | 2.3 kb | Michael G. Sargent |
| Sox10 | Ampicillin | pBSK | XhoI | T7 | EcoRI | T3 | 1.3 kb | JP. Saint-Jeannet |
| Snail2 | Ampicillin | pCS107 | | | EcoRI/BamHI | T7 | | EXRC |
| Xhe2 | Ampicillin | pBSK | | | XbaI | T7 | | AH.Monsoro-Burq |
| PAX6 | Ampicillin | pBSK | XhoI | T3 | XbaI | T7 | | Bill Harris |
| En2 | Ampicillin | pBSK | | | XbaI | T3 | 1.45 kb | Nancy Papalopulus |
| Zic1 | Ampicillin | | | | EcoRI | T3 | | Dr Jung Aruga |
| Sox2 | Ampicillin | pCS2+ | NotI | SP6 | EcoRI | T7 | | Prof. Yoshiki Sasai |

### Supplementary figure legends

**Supplementary Figure 1- MO and miRNA mimic validation experiments**. (A) q-RT-PCR data on miRNA expression levels in embryo following MO experiments. Error bars depict mean± s.e.m. (B) MiRNA sequences showing alignment of miRNA with mature miRNA, miRNA mimic used to rescue phenotypes, MO sequence, and see region of the miRNA. (C) miRNA mimic dose response, use lacZ 300 pg or GFP capped RNA as a tracer. No phenotypes were observed in the tadpoles. (D) q-RT-PCR validation on miRNA expression levels following miRNA mimic overexpression, miRNA KD, and then rescue using miRNA MO and miRNA mimic. Error bars depict mean± S.E.M with three biological and technical triplicates. Embryos were injected into both blastomeres at 2 cell stage. miR-219 rescue: miR-

219 mimic 15 µM vs miR-219 MO 60 ng p=0.008, miR-219 MO vs miR-219 MO 60 ng + miR-219 mimic 15 µM p=0.017, miR-219 mimic 15 µM vs miR-219 mimic 15 µM + miR-219 MO 60 ng p=0.039.

**Supplementary Figure 2 – miR-219 knockdown effect on NC marker *twist1***
MiR-219 knockdown in tailbud embryos (St. 27) perturbs *twist1* expression. Embryos were injected into one dorsal blastomere at 4-cell stage of development with MO and mismatch (MM) control. (A) WISH of st. 27 embryos with *twist1*. Lateral views of the injected and uninjected sides. The MO injected side clearly shows a decrease in *twist1* expression compared to the uninjected side and the MM control. (B) Count data for phenotype incidence. 3 biological replicates were obtained for mir-219 MO and MM miR-219. MO miR-219 n=229, MM miR-219 n=300.

**Supplementary Figure 3 – Snail2 and Pax3 MO phenotypes in Pax3 MO dose response**. Increasing doses perturbed Snail2 expression and reduced Pax3 with increasing dose up until 40 ng. Embryos were co-injected with 300 pg of lacZ cRNA tracer and developed with red-gal using a 10 nL calibrated needle. (A) Increasing doses of *Pax3* MO perturbed *Snail2* expression. (B) Increasing *Pax3* MO dose led to increasing loss of *Pax3* expression. *Pax3* MO based on published MO in: (Monsoro-Burq et al., 2005).

**Supplementary Figure 4- Cryosectioning of Pax3 phenotypes.** *X. laevis* embryos were co-injected with 300pg lacZ tracer, using a 10 nL calibrated needle. Injected side of the embryo is the right side. Embryos were injected at the 4-cell stage of development into 1 dorsal blastomere. (A) Embryos were injected with 40 ng of Pax3 MO. Whole mount view and cross-section showing loss of Pax3 in NC and NP regions. (B) Embryos were injected with 60 ng of miR-219 MO. Whole mount view and cross section showing expansion of Pax3 in superficial ectoderm and loss in NC area, Pax3 expression is lost in NC and NP regions. (C) Embryos were injected with 40 ng of Pax3 MO and 60 ng of miR-219 MO. (C) Whole-mount view and cross section showing a rescue of Pax3 expression in NC, NP and superficial ectoderm regions. Black lines through whole-mount views show the plane and region of the embryo sectioned. Sections imaged at 5x magnification. Black arrows indicate areas of phenotypic interest.

**Supplementary Figure 5 – Expression of the top 50 significantly differentially expressed gene following MO KD for miR-219.** Depleted genes are highlighted in blue with enriched genes in yellow.

**Supplementary Figure 6- Luciferase assay protocol.** Luciferase assay protocol used to assess EYA1 as a target of miR-219 Modified pGL3 plasmid with the region of the 3' UTR of EYA1 incorporated which includes the miR-219 binding site. Forward and Reverse EYA1 primers designed for mutagenesis are labelled in purple (A). A zoomed in version of the primers designed for mutagenesis of EYA1 3'UTR. By mutating three nucleotides the miR-219 target sequence was changed to a SalI restriction site (B). An alignment of the Eya1 3'UTR sequence of various organisms. The region highlighted in white in the target sequence for miR-219 (C). (D) A schematic drawing of the luciferase assay protocol. (D') Luciferase assay showing EYA1 is a target of miR-219 in vitro. The unmodified 3' UTR and the 3'UTR with the miR-219 target site mutated were assayed using chicken DF1cells with the miRNA mimics indicated below the graph. MiR-219 reduced luciferase activity by over 50% and this was rescued when the target site was mutated. For statistical analyses, a Mann-Whitney test was performed. For significance: $p < 0.05$: *; $p < 0.01$: **; p <0.001: ***; p <0.0001: ****

**Supplementary Figure 7- Assessment of miR-219 predicted target gene (Hes5.3 expression following MO mediated miRNA-KD.** Embryos were injected into the animal pole at the 2-cell stage of development with MO and 20 pg of GFP cRNA. Whole mount in situ hybridisation of *Hes5.3* (st. 14) following MO mediated miRNA KD. For miR-219 MM 60 ng n=39 (phenotype n=1) and for miR-219 MO 60 ng n=67 (phenotype n=44).

SFig. 1

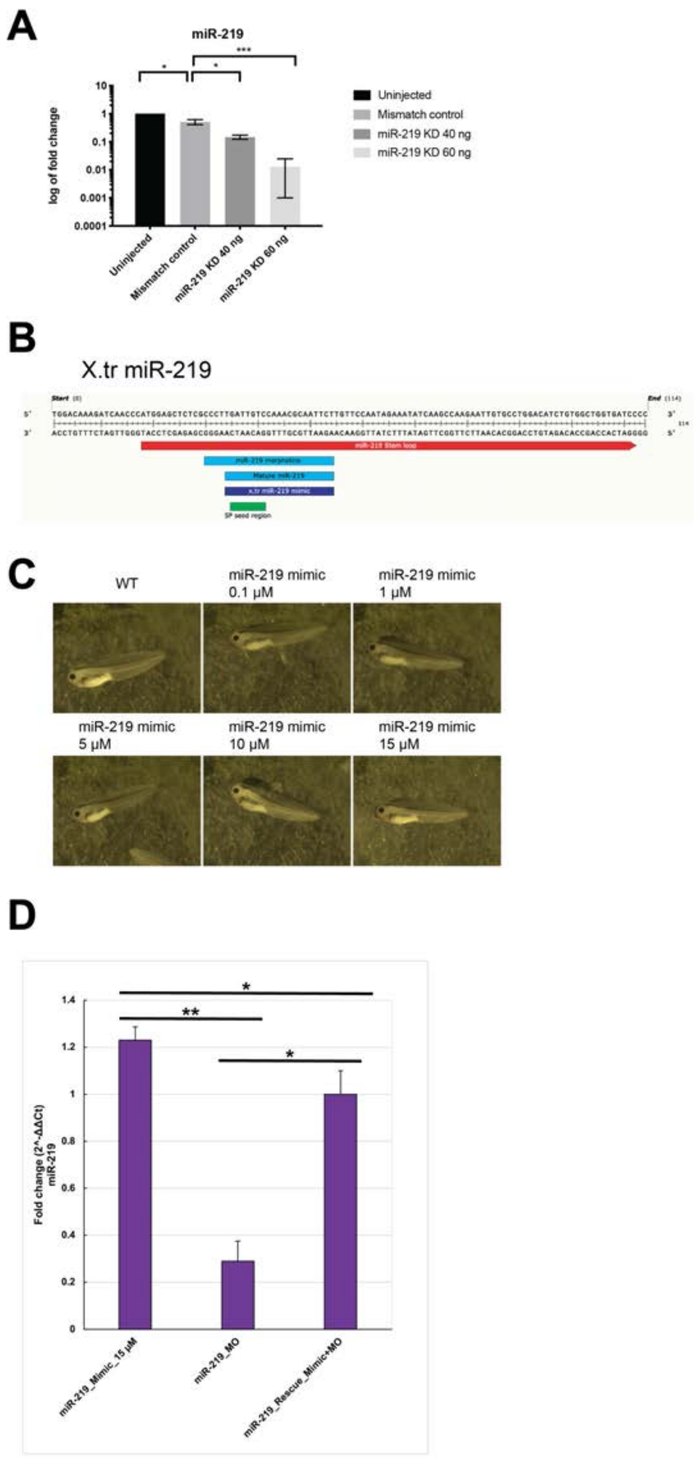
A
miR-219
log of fold change
10
1
0.1
0.01
0.001
0.0001
*
*
***
Uninjected
Mismatch control
miR-219 KD 40 ng
miR-219 KD 60 ng
B
X.tr miR-219
C
WT
miR-219 mimic 0.1 µM
miR-219 mimic 1 µM
miR-219 mimic 5 µM
miR-219 mimic 10 µM
miR-219 mimic 15 µM
D
Fold change (2^-ΔΔCt) miR-219
1.4
1.2
1
0.8
0.6
0.4
0.2
0
*
**
*
miR-219_Mimic_15 µM
miR-219_MO
miR-219_Rescue_Mimic+MO

SFig. 2

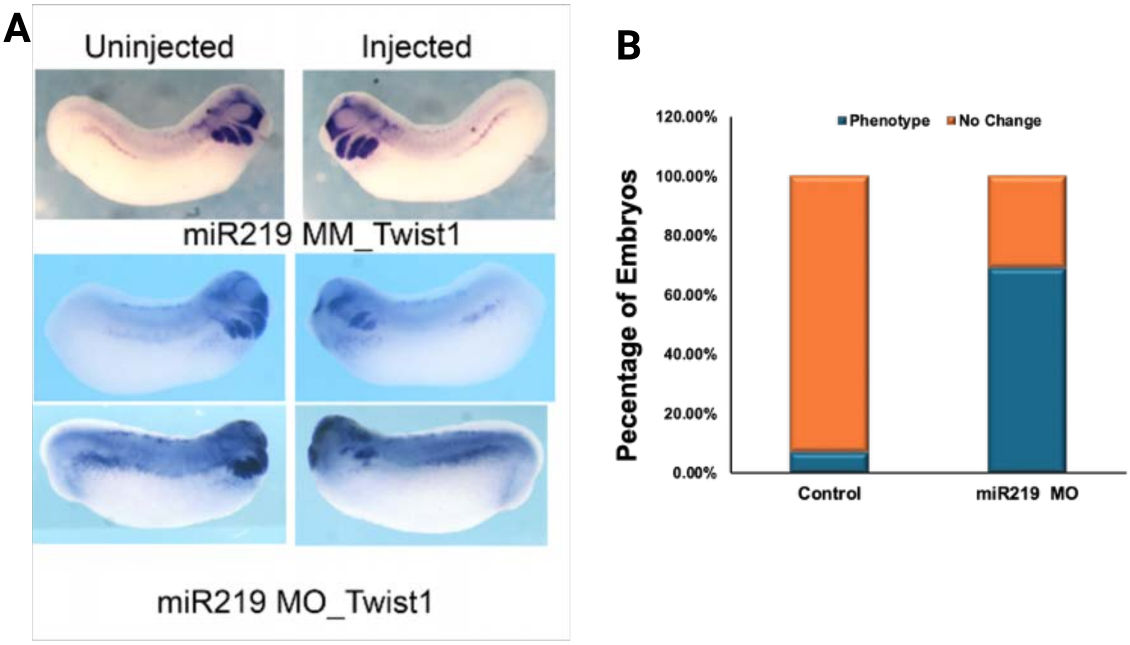
A
Uninjected
Injected
miR219 MM_Twist1
miR219 MO_Twist1
B
Phenotype
No Change
120.00%
100.00%
80.00%
60.00%
40.00%
20.00%
0.00%
Pecentage of Embryos
Control
miR219 MO

SFig. 3

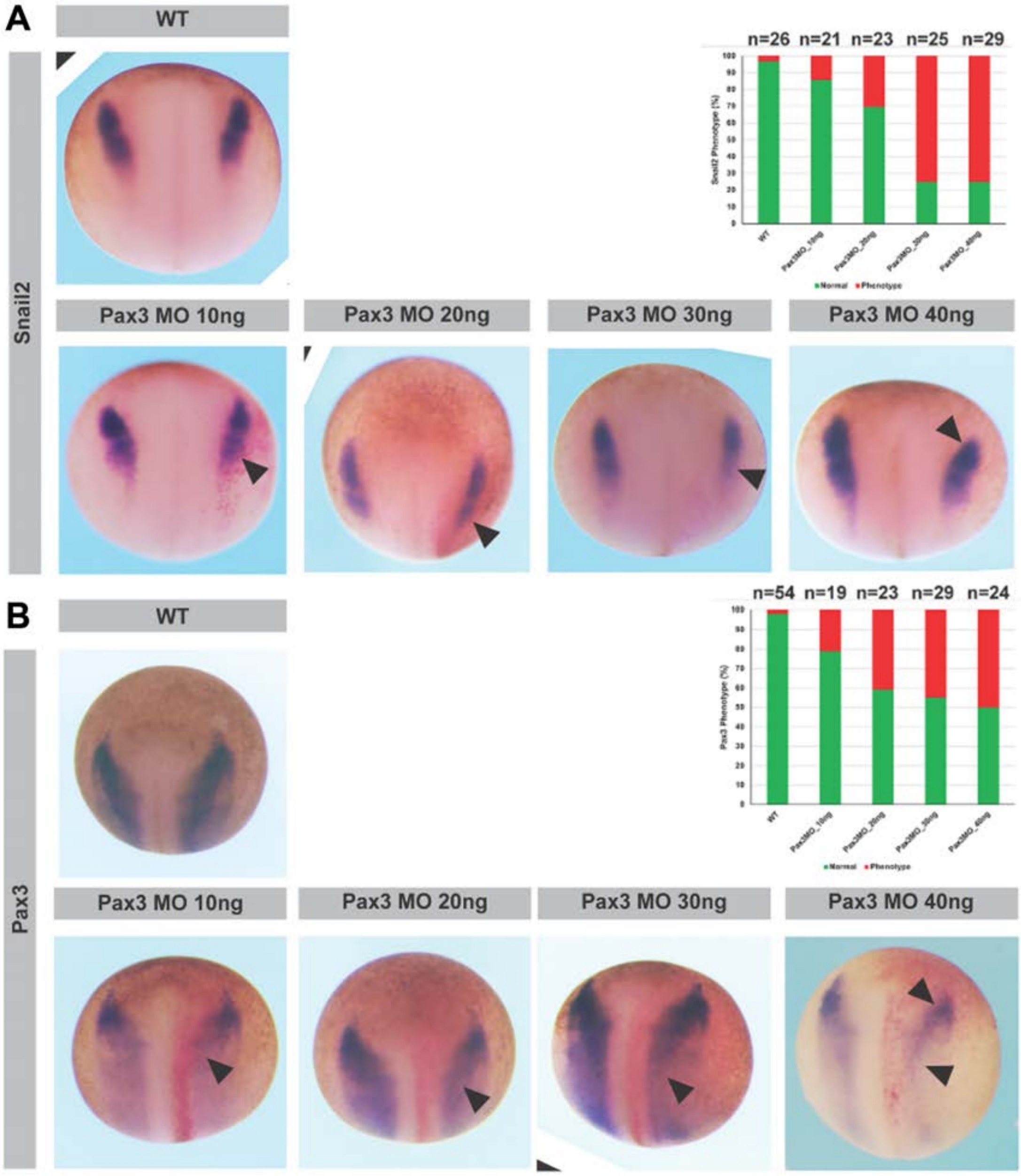
A
WT
n=26 n=21 n=23 n=25 n=29
Snail2
Pax3 MO 10ng
Pax3 MO 20ng
Pax3 MO 30ng
Pax3 MO 40ng
B
WT
n=54 n=19 n=23 n=29 n=24
Pax3
Pax3 MO 10ng
Pax3 MO 20ng
Pax3 MO 30ng
Pax3 MO 40ng

SFig. 4

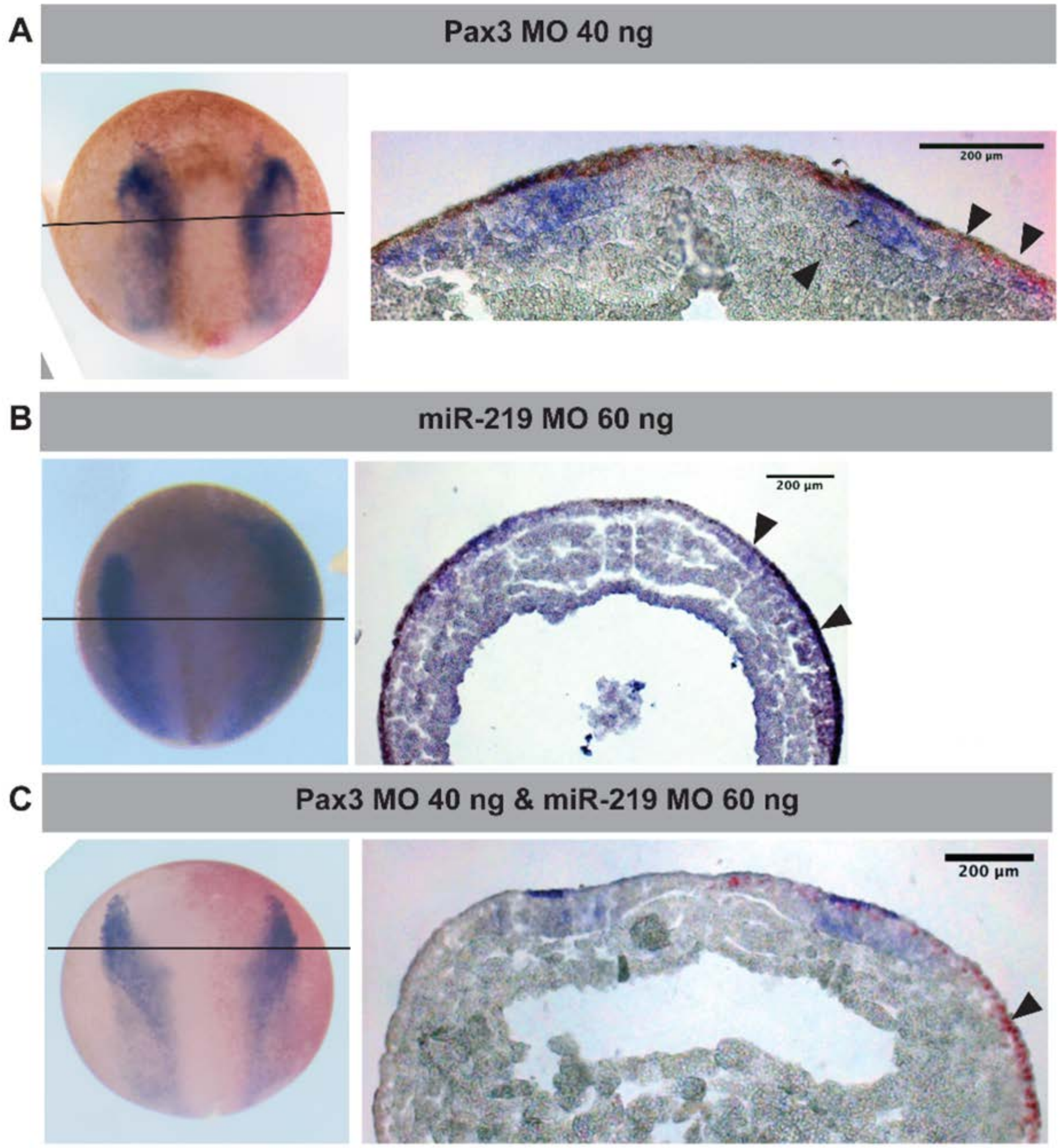
A
Pax3 MO 40 ng
200 µm
B
miR-219 MO 60 ng
200 µm
C
Pax3 MO 40 ng & miR-219 MO 60 ng
200 µm

SFig. 5

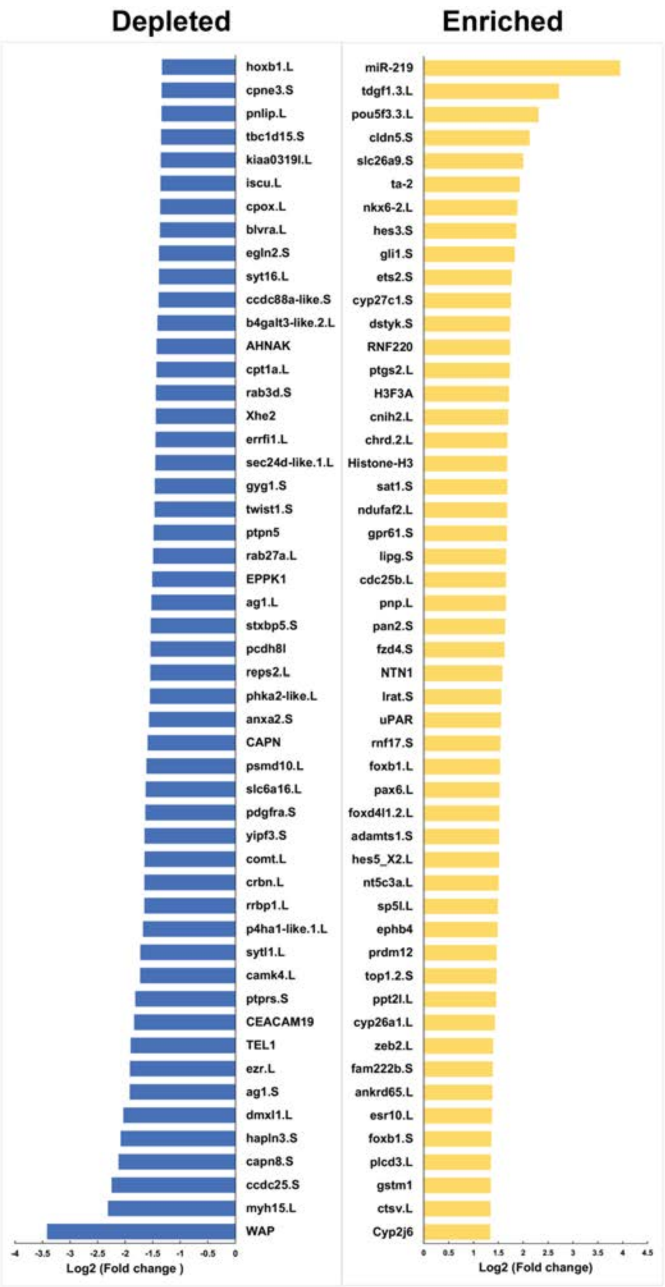
Depleted
Enriched
hoxb1.L
cpne3.S
pnlip.L
tbc1d15.S
kiaa0319l.L
iscu.L
cpox.L
blvra.L
egln2.S
syt16.L
ccdc88a-like.S
b4galt3-like.2.L
AHNAK
cpt1a.L
rab3d.S
Xhe2
errfi1.L
sec24d-like.1.L
gyg1.S
twist1.S
ptpn5
rab27a.L
EPPK1
ag1.L
stxbp5.S
pcdh8l
reps2.L
phka2-like.L
anxa2.S
CAPN
psmd10.L
slc6a16.L
pdgfra.S
yipf3.S
comt.L
crbn.L
rrbp1.L
p4ha1-like.1.L
sytl1.L
camk4.L
ptprs.S
CEACAM19
TEL1
ezr.L
ag1.S
dmxl1.L
hapln3.S
capn8.S
ccdc25.S
myh15.L
WAP
-4 -3.5 -3 -2.5 -2 -1.5 -1 -0.5 0
Log2 (Fold change )
miR-219
tdgf1.3.L
pou5f3.3.L
cldn5.S
slc26a9.S
ta-2
nkx6-2.L
hes3.S
gli1.S
ets2.S
cyp27c1.S
dstyk.S
RNF220
ptgs2.L
H3F3A
cnih2.L
chrd.2.L
Histone-H3
sat1.S
ndufaf2.L
gpr61.S
lipg.S
cdc25b.L
pnp.L
pan2.S
fzd4.S
NTN1
lrat.S
uPAR
rnf17.S
foxb1.L
pax6.L
foxd4l1.2.L
adamts1.S
hes5_X2.L
nt5c3a.L
sp5l.L
ephb4
prdm12
top1.2.S
ppt2l.L
cyp26a1.L
zeb2.L
fam222b.S
ankrd65.L
esr10.L
foxb1.S
plcd3.L
gstm1
ctsv.L
Cyp2j6
0 0.5 1 1.5 2 2.5 3 3.5 4 4.5
Log2 (Fold change)

SFig. 6

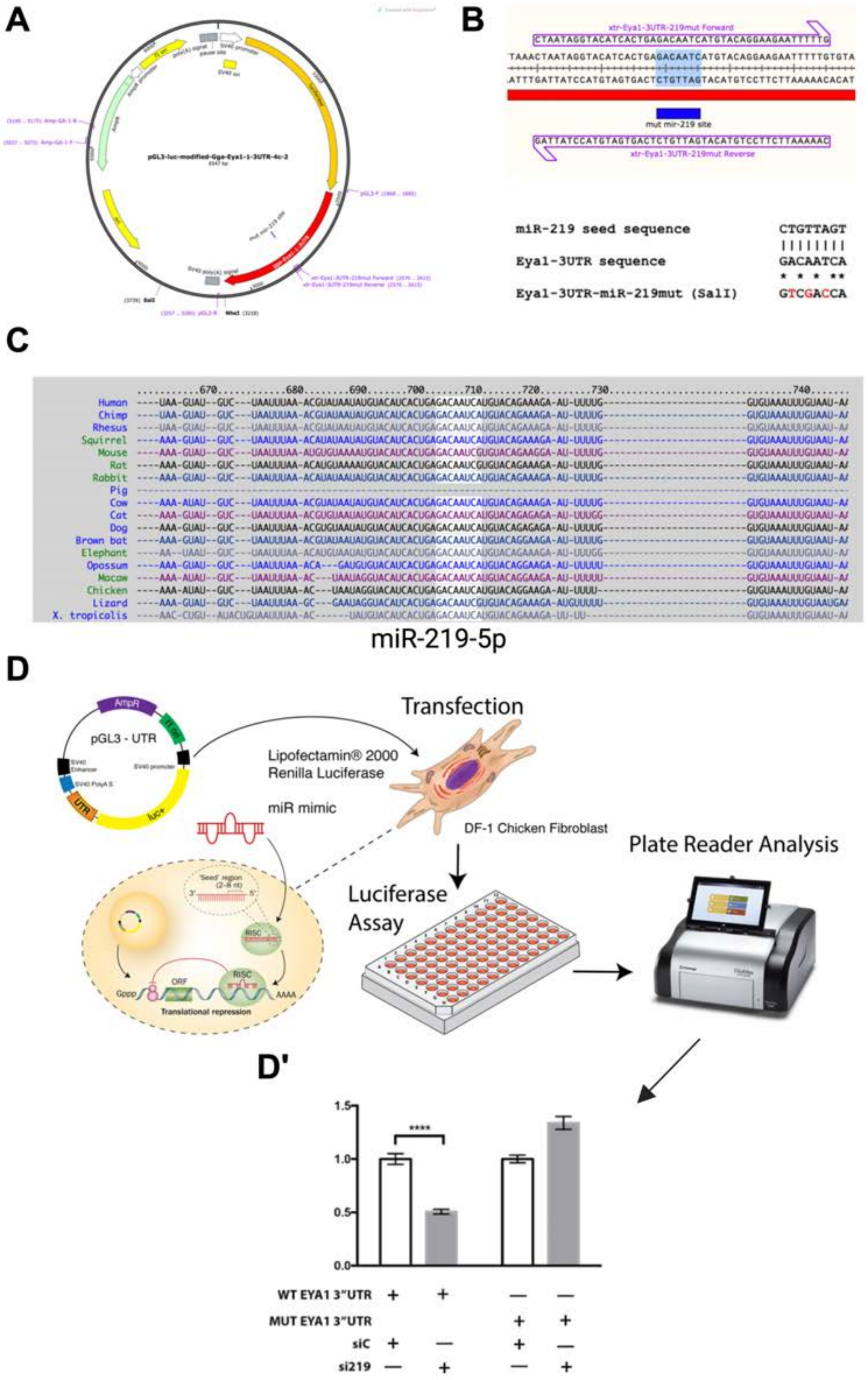
A
B
xtr-Eya1-3UTR-219mut Forward
mut mir-219 site
xtr-Eya1-3UTR-219mut Reverse
miR-219 seed sequence
CTGTTAGT
Eya1-3UTR sequence
GACAATCA
Eya1-3UTR-miR-219mut (SalI)
GTCGACCA
C
Human
Chimp
Rhesus
Squirrel
Mouse
Rat
Rabbit
Pig
Cow
Cat
Dog
Brown bat
Elephant
Opossum
Macaw
Chicken
Lizard
X. tropicalis
miR-219-5p
D
AmpR
pGL3 - UTR
SV40 promoter
UTR
luc+
Transfection
Lipofectamin® 2000
Renilla Luciferase
miR mimic
DF-1 Chicken Fibroblast
Plate Reader Analysis
Luciferase
Assay
ORF
RISC
AAAA
Translational repression
D'
1.5
1.0
0.5
0.0
****
WT EYA1 3"UTR
MUT EYA1 3"UTR
siC
si219

SFig. 7

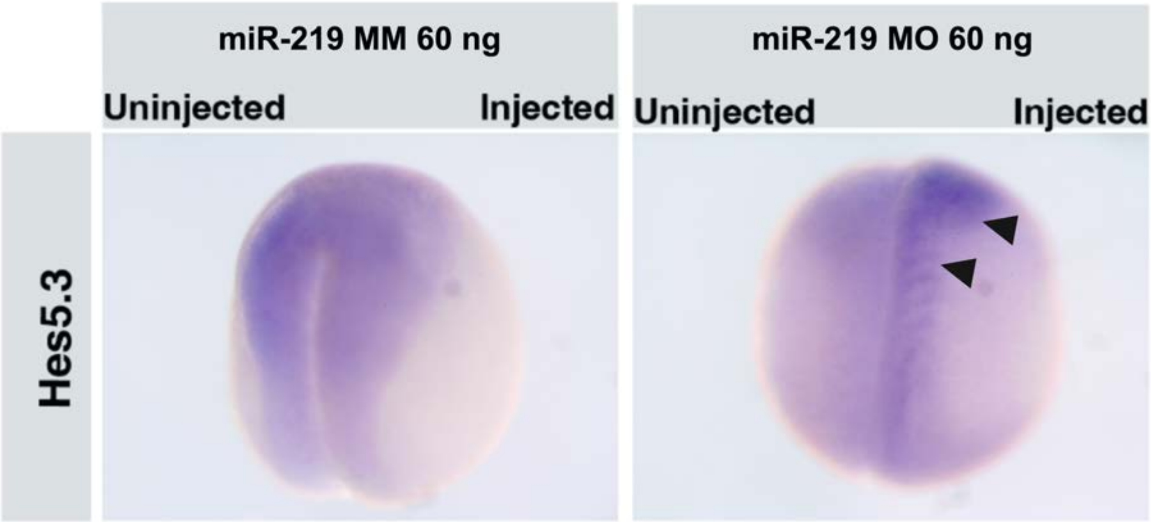

miR-219 MM 60 ng
miR-219 MO 60 ng
Uninjected
Injected
Uninjected
Injected
Hes5.3
